# Adaptive Reduction of Male Gamete Number in a Selfing Species

**DOI:** 10.1101/272757

**Authors:** Takashi Tsuchimatsu, Hiroyuki Kakui, Misako Yamazaki, Cindy Marona, Hiroki Tsutsui, Afif Hedhly, Dazhe Meng, Yutaka Sato, Thomas Städler, Ueli Grossniklaus, Masahiro M. Kanaoka, Michael Lenhard, Magnus Nordborg, Kentaro K. Shimizu

**Affiliations:** Department of Evolutionary Biology and Environmental Studies, University of Zurich, 8057 Zurich, Switzerland; Department of Plant and Microbial Biology & Zurich-Basel Plant Science Center, University of Zurich, 8008 Zurich, Switzerland; Gregor Mendel Institute, Austrian Academy of Sciences, A-1030 Vienna, Austria; Department of Biology, Chiba University, Chiba 263-8522, Japan; Kihara Institute of Biological Research, Yokohama City University, Yokohama 244-0813, Japan; Institute of Biochemistry and Biology, University of Potsdam, 14476 Potsdam, Germany; Graduate School of Science, Nagoya University, Nagoya 464-8602, Japan; JST ERATO Higashiyama Live-Holonics Project, Nagoya University, Nagoya 464-8602, Japan; Graduate School of Bioagricultural Sciences, Nagoya University, Nagoya 464-8601, Japan; Molecular and Computational Biology, University of Southern California, Los Angeles, California, 90089-0371 USA; Institute of Integrative Biology, ETH Zurich, 8092 Zurich, Switzerland

## Abstract

The number of male gametes produced is critical for reproductive success and varies greatly between and within species^1–3^. Evolutionary reduction of male gamete production has been widely reported in plants as a hallmark of the selfing syndrome, as well as in humans. Such a reduction may simply represent deleterious decay^4–7^, but evolutionary theory predicts that breeding systems could act as a major selective force on male gamete number: while large numbers of sperm should be produced in highly promiscuous species because of male–male gamete competition^1^, reduced sperm numbers may be advantageous at lower outcrossing rates because of the cost of gamete production. Here we used genome-wide association study (GWAS) to show a signature of polygenic selection on pollen number in the predominantly selfing plant *Arabidopsis thaliana*. The top associations with pollen number were significantly more strongly enriched for signatures of selection than those for ovule number and 107 phenotypes analyzed previously, indicating polygenic selection^8^. Underlying the strongest association, responsible for 20% of total pollen number variation, we identified the gene *REDUCED POLLEN NUMBER 1* affecting cell proliferation in the male germ line. We validated its subtle but causal allelic effects using a quantitative complementation test with CRISPR-Cas9-generated null mutants in a nonstandard wild accession. Our results support polygenic adaptation underlying reduced male gamete numbers.

---

Pollen number in seed plants and sperm number in animals have been studied extensively from agricultural, medical, and evolutionary viewpoints^3^. For example, the number of pollen grains decreased during rice domestication^9^. Further, it has been suggested that sperm number and quality may have declined in humans during the twentieth century. In flowering plants, evolutionary shifts from outcrossing to selfing through loss of self-incompatibility have occurred frequently, and selfing species generally show lower pollen numbers, a hallmark of the “selfing syndrome”^2,10–12^. A reduction in the number of male gametes could be caused by environmental changes, by the accumulation of mutations because of relaxed constraints, or by inbreeding depression, given the prevalence of male sterility in inbred individuals^4–7^. Alternatively, it may reflect adaptive evolution in response to the change in breeding system^2^.

A major hindrance in studying the mechanisms and evolution of the regulation of male gamete number is the almost unknown molecular basis of this quantitative trait. Genome-wide association studies (GWAS) provide an alternative approach to traditional mutant screens, exploiting natural genetic variation and recombination within species. Although there have been several GWAS investigating genes affecting gamete number, there is little experimental support for the validation of identified candidate loci^13^. Here we focus on the predominantly selfing plant *Arabidopsis thaliana*^2,14^. To determine variability in pollen number on a species-wide scale, we examined pollen number per flower for 144 natural accessions (Fig. 1a–d; Extended Data Tables 1 and 2) and found approximately four-fold variation (Fig. 1e). The consistency of this variation was validated using histological sections of stamens from representative accessions (Fig. 1c, d). We also measured the number of ovules per flower but did not find significant correlations between the number of male and female gametes (Extended Data Table 3), although negative correlations have often been reported in between-species comparisons, and are expected on theoretical grounds due to trade-offs in resource allocation^15,16^. Furthermore, we found that pollen number per flower was not significantly correlated with any of the 107 published phenotypes of flowering, defence-related, ionomic, and developmental traits (Extended Data Table 4; Supplementary Note)^17^, nor with climate variables or *S*-haplogroups (Extended Data Table 5)^18,19^, suggesting that variation in pollen number is largely independent of other traits.

**Figure 1.**
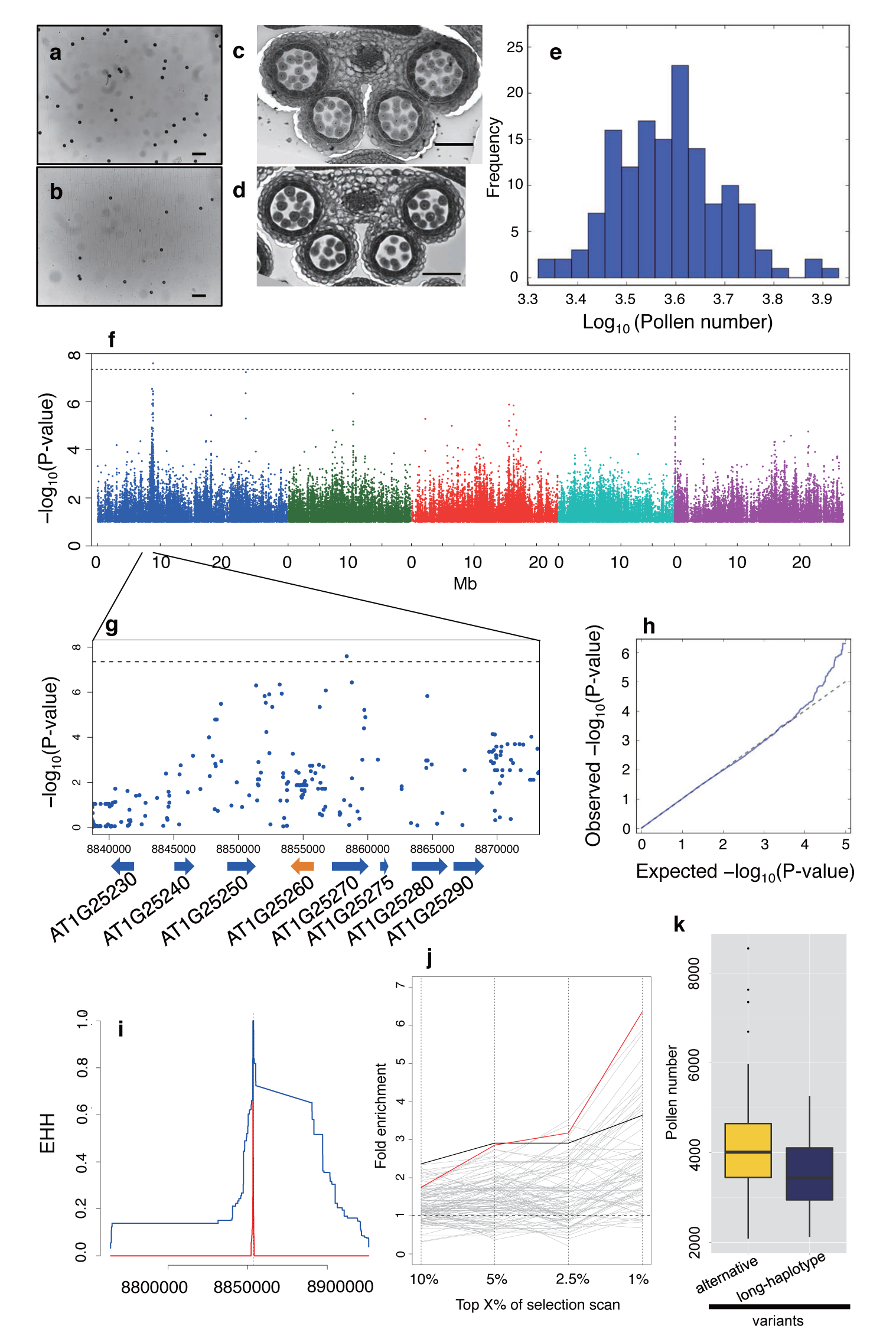
Genome-wide association studies (GWAS) of pollen number variation in *Arabidopsis thaliana*. (**a, b**) Pollen grains of Bor-4 (**a**) and Mz-0 (**b**) mounted on glass slides for counting^15^. Scale bars = 100 µm. (**c, d**) Histological sections of Bor-4 (**c**) and Mz-0 (**d**) stamens. Scale bars = 50 µm. **e,** Pollen number variation of 144 natural accessions. **f**, Manhattan plot of the GWAS. **g**, Closer view of the region around the significant GWAS peak on chromosome 1 with gene models and coordinates. (**f** and **g**) SNPs with minor allele frequency > 0.15 are shown; horizontal dashed lines indicate the nominal *P* < 0.05 threshold after Bonferroni correction. **h**, Quantile–quantile plot of the GWAS. **i**, Extended haplotype homozygosity (EHH) detected in the *RDP1* genomic region. Blue and orange lines correspond to the long haplotype and alternative variants, respectively. **j**, Signatures of selection at pollen number-associated loci. Each line indicates a phenotype (red: pollen number, black: ovule number, gray: 107 phenotypes^17^). The x-axis quantifies the extreme tails of the iHS statistic. The pollen and ovule GWAS show significant enrichment (permutation test, *P* < 0.05 cutoff for all iHS statistical tails; Extended Data Fig. 2). **k**, Accessions with the long-haplotype variant (defined by SNP “1-8853454”) generally showed lower pollen number (*P* = 0.000343, *t*-test; population structure-corrected GWAS *P* = 0.00588).

To evaluate genome-wide signatures of natural selection on gamete numbers, we first performed GWAS for pollen and ovule numbers using an imputed single nucleotide polymorphism (SNP) dataset for these lines (Fig. 1f, h; Extended Data Fig. 1). GWAS identified multiple peaks of association for pollen and ovule numbers, suggesting the polygenic nature of these traits. Focusing on the identified GWAS peaks, we performed an enrichment analysis to ask whether pollen and ovule number-associated peaks are enriched in long-haplotype regions, which could be due to partial or ongoing sweeps of segregating polymorphisms^20–22^. Using a haplotype-based iHS selection scan, we found that 10-kb windows including pollen number-associated loci (68 regions in total with *P* < 10^−4^) were significantly enriched in extreme iHS tails (*P* < 0.05, permutation test; Fig. 1i, j; Extended Data Fig. 2). Similarly, 10-kb windows including ovule number-associated loci also showed enrichment, albeit less than for pollen number (Fig. 1j). To assess the possibility that GWAS peaks could be associated with high iHS regions by chance, we compared these results with the GWAS of 107 phenotypes. We found that the iHS enrichment of the pollen number GWAS (*P* = 0.002 for the top 1% iHS tail; Extended Data Fig. 2) was among the highest, compared with many known adaptive traits included in the 107 phenotypes such as leaf number at flowering time and resistance to *Pseudomonas* pathogens^17,22^ (Extended Data Table 6). In addition, iHS enrichment of the ovule number GWAS was also significant (*P* = 0.030 for the top 1% iHS tail; Extended Data Fig. 2). This enrichment supports polygenic selection on male and female gamete numbers at a considerable number of loci throughout the genome.

**Figure 2.**
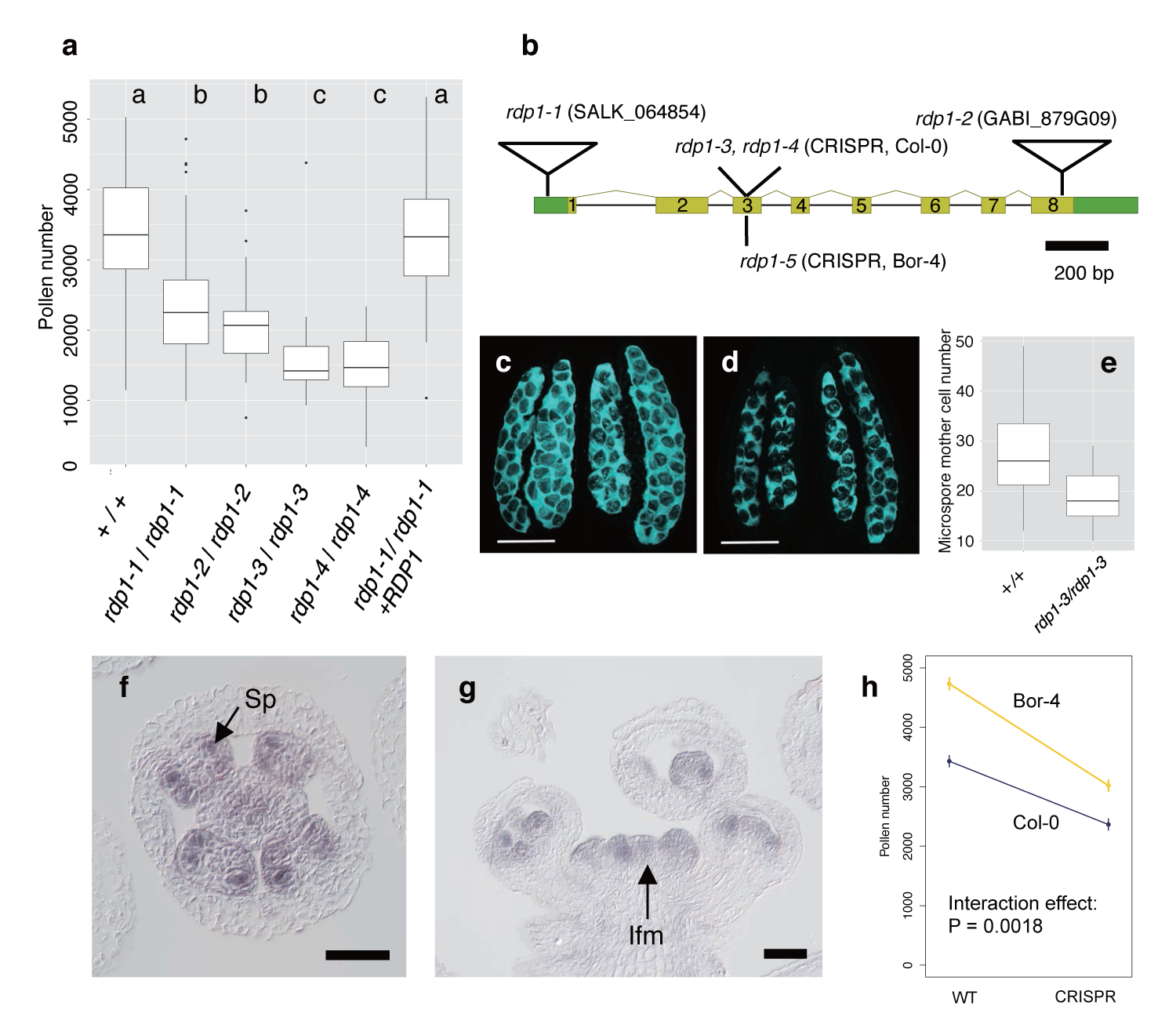
Functional characterization of the *RDP1* gene. **a**, Pollen number differences between four homozygous mutants (*rdp1-1/rdp1-1*, N = 132; *rdp1-2/rdp1-2*, N = 45; *rdp1-3/rdp1-3*, N = 40; and *rdp1-4/rdp1-4*, N = 60), wild type (+/+, N = 135), and the complemented lines with the Col-0 allele (*rdp1-1*/*rdp1-1*+*RDP1*, N = 411). Letters (a, b, c) indicate significant differences in pollen number, determined by nested analysis of variance and *post hoc* Tukey test; *P* < 0.05. **b**, Schematic structure of the *RDP1* gene. Untranslated regions (green boxes), exons (yellow boxes), introns (bars), two T-DNA insertion mutants (*rdp1-1, rdp1-2*; open triangles), and three CRISPR frameshift mutants (*rdp1-3* and *rdp1-4* on Col-0, and *rdp1-5* on Bor-4 backgrounds, respectively) are shown. (**c**, **d**) Microspore mother cells of +/+ (C) and *rdp1-3/rdp1-3* (D) stained with aniline blue. Dark spots indicate microspore mother cells. Scale bars: 50 µm. **e**, The number of microspore mother cells was significantly lower in the homozygous mutant *rdp1-3/rdp1-3* (N = 40) than in the wild type (N = 50, Wilcoxon rank sum test, *P* = 2.82e–06). (**f**, **g**) *In situ* hybridization for *RDP1*. **f**, Stage 8 floral cross section; expression is detected in sporogenous cells (Sp). **g**, Inflorescence longitudinal section; hybridization signal is detected in the inflorescence meristem (Ifm) and young flowers. Scale bars: 50 µm. **h**, Pollen number of wild-type and null mutants generated using the CRISPR-Cas9 technique in Col-0 (blue) and Bor-4 (yellow) accessions. Dots and error bars indicate means and standard errors, respectively. The mutation in Bor-4 had larger effects on pollen number than that in Col-0 (ANOVA interaction effect *P* = 0.0018).

To further examine the nature of the putative targets of selection, we tried to identify the genes underlying pollen number variation; however, the GWAS for pollen number did not yield hits for genes with known function in early stamen or pollen development in the vicinity of any of the top five peaks. To obtain experimental evidence concerning genes underlying pollen number variation, we conducted functional analyses of the genes in the region of the highest peak of the pollen number GWAS, which explains about 20% of the total phenotypic variance and satisfies the criterion for genome-wide significance (–log_10_ *P* = 7.60). This region is of particular interest because it also satisfies the criteria for genome-wide significance of the iHS statistic (*P* = 0.0149; Fig. 1i, Extended Data Fig. 3), suggesting a selective sweep. To test whether the signature of selection in this region might be due to other traits, we examined whether there is an association signal for any of the 107 published phenotypes, ovule number, or variants showing climatic correlations^17,18^. In the 10-kb window including the SNP of the highest GWAS score for pollen number, we found no genotype–phenotype associations below *P* < 10^−5^ or climate– SNP correlations below an empirical *P* < 0.01, i.e., there is no evidence for selection on traits other than pollen number. We also found that accessions with the long-haplotype variant produced lower pollen numbers than those with alternative haplotype variants (*P* = 0.000343, *t*-test; population structure-corrected GWAS *P* = 0.00588; Fig. 1k), indicating selection for reduced pollen number.

**Figure 3.**
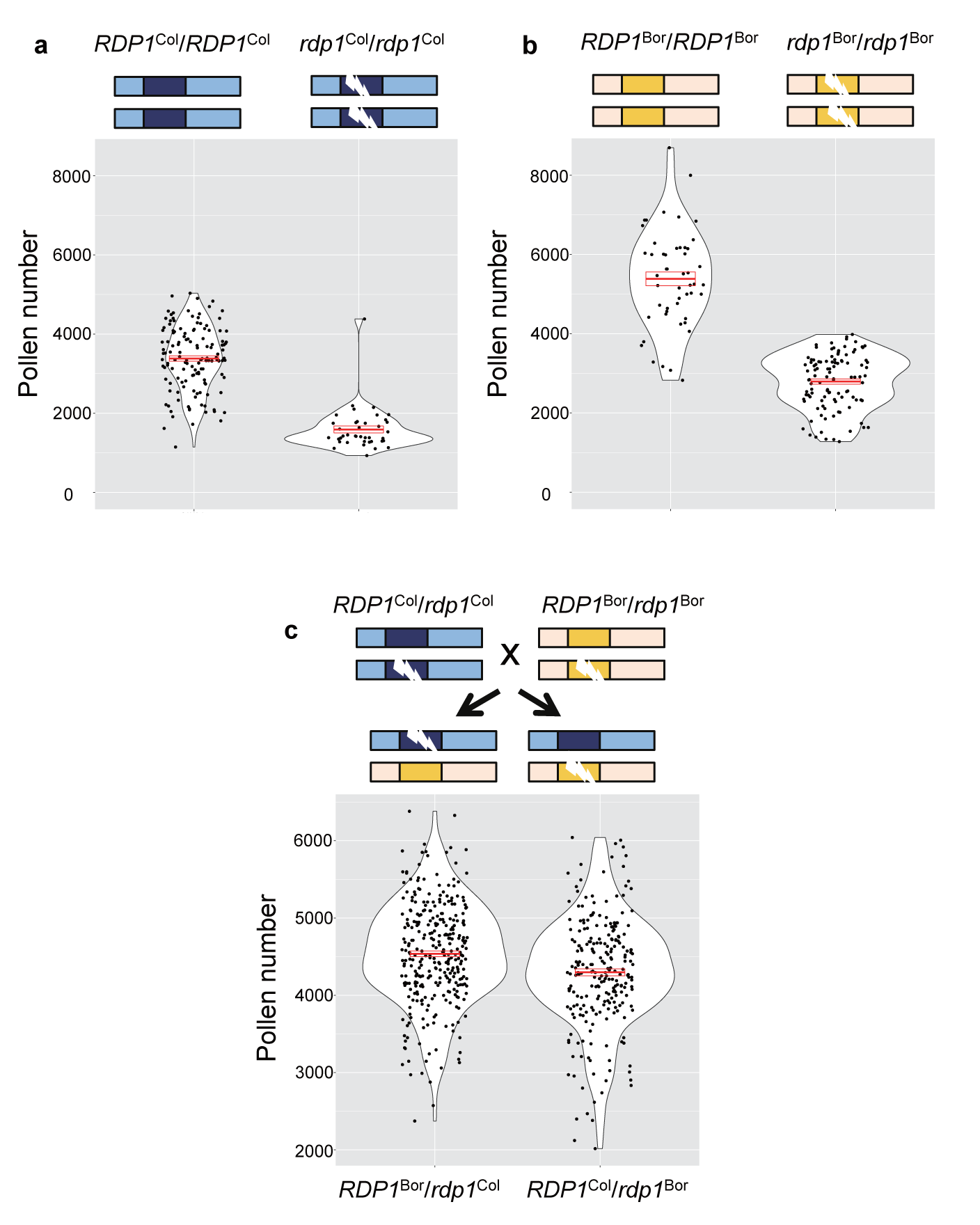
Quantitative complementation test of the *RDP1* gene. Violin plots with means and standard errors indicated by red bold bars and boxes, respectively. (**a**, **b**) Pollen number difference between wild-type and a homozygote of a frameshift allele generated by the CRISPR-Cas9 technique in the Col-0 background (**a**, *RDP1*^*Col*^*/RDP1*^*Col*^, N = 135; *rdp1*^*Col*^*/rdp1*^*Col*^, N = 40) and in the Bor-4 background (**b**, *RDP1*^*Bor*^*/RDP1*^*Bor*^, N = 50; *rdp1*^*Bor*^*/rdp1*^*Bor*^, N = 106). **c**, The difference in the effect on pollen number by two natural alleles, *RDP1*^*Col*^ and *RDP1*^*Bor*^. Pollen number of plants with *RDP1*^*Col*^ was significantly less than that of plants with *RDP1*^*Bor*^ (nested analysis of variance; *P* = 1.10 × 10^−5^; *rdp1*^*Col*^*/RDP1*^*Bor*^, N = 315; *RDP1*^*Col*^*/rdp1*^*Bor*^, N = 247). The two alleles were compared in heterozygous state with a frameshift CRISPR-Cas9 allele, with otherwise identical genomic backgrounds. F_1_ plants were obtained from the cross of two heterozygotes, *RDP1*^*Col*^*/rdp1*^*Col*^and *RDP1*^*Bor*^*/rdp1*^*Bor*^.

Of the three genes in this chromosomal region with the highest GWAS scores (*AT1G25250, AT1G25260*, and *AT1G25270*; Fig. 1g), the expression level of *AT1G25260*, a gene of unknown function, was much higher in flower buds than that of the other two genes (Extended Data Fig. 4). We therefore analyzed pollen numbers in mutants of *AT1G25260* and identified two T-DNA insertion mutants that showed a 32% reduction in pollen number (Fig. 2a). We hereafter refer to *AT1G25260* as *REDUCED POLLEN NUMBER 1* (*RDP1*). Because both *rdp1-1* (insertion in the 5´ UTR) and *rdp1-2* (insertion at the end of the coding sequence) (Fig. 2b) still showed low levels of expression, they are likely to be hypomorphic mutants (Extended Data Fig. 4). We generated two amorphic (null) frameshift mutants of *RDP1* (*rdp1-3* and *rdp1-4*) using the CRISPR-Cas9 system^23,24^; these indeed showed an even greater reduction in pollen number, but still produced about half the number of pollen grains of the wild type (53% for *rdp1-3*; Fig. 2a), supporting the quantitative nature of the effect of *RDP1*. Pollen size was slightly increased, consistent with the well-known negative relationship between pollen number and size, even within the same genotype (Extended Data Fig. 5)^15^. The mutant phenotype was complemented by transforming a 4.3-kb genomic fragment of the Col-0 accession encompassing *RDP1* (Fig. 2a, Extended Data Fig. 5). In contrast, an insertion into the neighbouring gene *AT1G25250* almost completely abolished its expression but did not result in any significant change in pollen number (Extended Data Fig. 6). The phenotype of four independent mutants together with the successful complementation test thus demonstrated that *RDP1* is involved in the control of pollen number per flower.

Based on phylogenetic analysis, RDP1 is a putative homolog of yeast mRNA turnover protein 4 (Mrt4) (Extended Data Figs 7 and 8). The Mrt4 gene is nonessential in yeast, and its null mutant shows a phenotype of slightly slower growth. The Mrt4 protein shares similarity with the ribosome P0 protein and is necessary for the assembly of the P0 protein into the ribosome. The human ribosome P0 gene is reported to have an extra-ribosomal function in cancer through modulating cell proliferation^25,26^. During wild-type anther development, sporogenous cells first divide and differentiate into microspore mother cells (microsporocytes)^27^. Then, the mature pollen grains containing the male gametes are formed through meiosis and two mitotic divisions. The null *rdp1*-*3* mutant produced fewer microspore mother cells than the wild type (Fig. 2c–e), indicating a reduction in cell numbers before meiosis. Consistent with this, *in situ* mRNA hybridization experiments detected strong expression of *RDP1* in sporogenous and microspore mother cells, but not in microspores (Fig. 2f, g; Extended Data Fig. 9). *RDP1* was also expressed in other proliferating cells, including those in inflorescence and floral meristematic regions and ovule and petal primordia (Extended Data Figs 4 and 9), supporting a more widespread role in proliferating cell types with putatively high demands for ribosome biogenesis^28^.

Fusing the *RDP1* promoter to the *GUS* reporter gene to assess its activity confirmed the expression pattern in stamens (Extended Data Fig. 10) and demonstrated marked expression in root tips and young leaf primordia during the vegetative phase; these data are supported by real-time PCR (Extended Data Fig. 4b). Consistent with *RDP1* expression in proliferating tissues, the *rdp1-3* null mutant showed pleiotropic phenotypes, including slower vegetative growth and reduced ovule numbers per flower (Extended Data Fig. 11). Because these pleiotropic phenotypes would be deleterious in natural environments, we hypothesize that natural variants of *RDP1* are not null mutations. In summary, these data suggest that *RDP1* is expressed and required in proliferating *A. thaliana* cells, yet the natural variants we have identified predominantly affect proliferation of sporogenous cells in the anthers, consistent with the function of its yeast homologue in cell proliferation.

It has been difficult to experimentally detect subtle allelic effects of single loci on quantitative traits^29^. When allelic effects are subtle, transgenic analysis of natural alleles is not sufficiently powerful because the phenotypes of transgenic *A. thaliana* plants tend to be highly variable as a result of experimental uncertainties such as the transgene insertion sites. In contrast, a complementation test can determine the effects of natural alleles in heterozygous state with a null allele if the effects of other loci are small, although this may be confounded by polygenic effects in the genomic background. To conduct quantitative complementation^29,30^, we took advantage of the CRISPR-Cas9 technique to generate frameshift null alleles in a nonstandard natural accession, Bor-4, which has a high-pollen-number phenotype, in addition to the standard Col-0 accession (Fig. 2b; Fig. 3a–c). In both accessions, *rdp1* CRISPR null mutants showed reduced pollen number (*P* = 9.41 × 10^−14^ for Col-0, *P* < 2.0 × 10^−16^ for Bor-4; Fig. 3a, b; Fig. 2h), supporting the idea that naturally occurring variants are not null. Disruption of *RDP1* had a stronger effect on pollen number in Bor-4 than in Col-0 (ANOVA interaction effect *P* = 0.0018; Fig. 2h). This finding supports the notion that the Bor-4 allele has a larger effect on pollen number than the Col-0 allele, although other loci in the genetic backgrounds of Bor-4 and Col-0 may contribute to this difference through epistasis.

To exclude a potential epistatic contribution, we utilized a quantitative complementation test that controls for the genetic background (Fig. 3). Among the F_1_ plants obtained by crossing heterozygotes of each genetic background, we compared two genotypes: *RDP1*^*Col*^*/rdp1*^*Bor*^ vs. *RDP1*^*Bor*^*/rdp1*^*Col*^. These lines are genetically identical except for the differences at *RDP1*, where they both carry a frameshift allele but differ with respect to the functional allele; because of the crossing design, any independently segregating off-target effects would be equally distributed between the two genotype cohorts of interest.We found that pollen number in plants with *RDP1*^*Col*^ was significantly lower than in plants with *RDP1*^*Bor*^ (*P* = 1.10 × 10^−5^; Fig. 3c), providing evidence for causal allelic differences at *RDP1*.

To our knowledge, *RDP1* is the first gene shown to underlie natural variation in male gamete numbers, and our study provides evidence for polygenic selection on pollen number-associated loci, including *RDP1*. Even though *RDP1* encodes a ribosome-biogenesis factor that would be required globally for proliferative growth, the naturally selected alleles predominantly confer reduced pollen number, a hallmark of the selfing syndrome. This is analogous to a hypomorphic allele of the human *G6PD* gene, which encodes an enzyme in the pentose phosphate pathway and features a long haplotype because of selection for malaria resistance^21^. Our work illustrates that a combination of GWAS and functional analysis performed with a quantitative complementation test based on the CRISPR-Cas9 technique provides a powerful approach to study the evolution of quantitative natural variation in any species.

## Methods Summary

### Genome-wide association study (GWAS)

GWAS was performed to identify loci associated with pollen number variation in the 144 natural accessions. The median values of pollen number were used to represent each accession. We used the log-transformed values of pollen number for GWAS, as they did not deviate significantly from a normal distribution (Shapiro–Wilk normality test: *P* = 0.438). We used the mixed model, in which a genome-wide kinship matrix was incorporated as a random effect^31^.

### *In situ* hybridization

Flower buds were fixed with FAA and dehydrated through an ethanol series. Fixed samples were embedded in paraplast. *RDP1* cDNA was amplified using the primer pair (At1g25260g2F; 5´-tgcctaatcaaagcgagtagacc-3´ and At1g25260gR; 5´-cagagcaagttcagcttgaaagtagc-3´). Cloned cDNA was used as a template for *in vitro* transcription using a MAXIscript T7 labeling kit (Thermo). *In situ* mRNA hybridization was performed as described previously^32^.

### CRISPR mutants

To generate *RDP1* frameshift mutants, the CRISPR-Cas9 system was used as previously described^24^. A 20-nucleotide target sequence (5´-GCAGAAGATGAACTCCGTTC-3´) was selected. For plant transformation, the binary vector was first introduced into *Agrobacterium tumefaciens* (GV3101) and then into *A. thaliana* by the floral-dip method.

### Selection scan

We used the iHS statistic for the selection scan; this statistic compares the extended haplotype homozygosity (EHH) between two alleles by controlling for the allele frequency of each SNP^20^. We asked whether the GWAS-associated 10-kb genomic windows were enriched significantly in the extreme tails of the iHS statistic.

## Supplementary Information

is available in the online version of the paper.

### Acknowledgments

We thank Matt Horton, Robert Martin, Atsushi Tajima and Hirokazu Tsukaya for discussion and the Functional Genomics Center Zurich for technical support. This study was supported by the Swiss National Science Foundation and JST CREST Grant Number JPMJCR16O3, Japan, to KKS, a PSC-Syngenta fellowship to TT, TS, and KKS, the University Research Priority Program of Evolution in Action of the University of Zurich to KKS and UG, MEXT KAKENHI Grant Numbers 16H06469, 16K21727, 26113709 to KKS, 16H06467, 17H05833 to TT and 16H06465 to MK, the Forschungskredit of the University of Zurich and an EMBO long-term fellowship to TT, a JSPS postdoctoral fellowship for research abroad to HK and TT, and an Advanced Grant of the European Research Council to UG.

## Author Contributions

T.T., H.K., M.Y., T.S., M.L., M.N. and K.K.S. conceived and designed the study; T.T., H.K., M.Y., D.M. and K.K.S analyzed the data with help from T.S., M.L. and M.N; T.T., H.K., M.Y., C.M., H.T., A.H., Y.S. and M.K. performed experiments and generated the data; T.T., H.K., M.Y. and K.K.S. wrote the paper with inputs from U.G., T.S., M.L. and M.N; All authors discussed the results and commented on the manuscript.

## Author Information

Sequence data have been deposited GenBank under accession numbers LC164158-LC164163. The authors declare no competing financial interests. Correspondence and requests for materials should be addressed to K.K.S.

## Methods

### Pollen and ovule counting for genome-wide association studies (GWAS)

To perform GWAS, numbers of pollen grains and ovules per flower were counted for the 144 and 151 worldwide natural accessions, respectively (Extended Data Tables 1–3). Plants were grown at 21 °C under a 16 h light/8 h dark cycle without vernalization. We grew four plants per accession. Three flower buds per plant were harvested from the main inflorescence, and each flower bud was collected into a 1.5 ml tube and dried at 65 °C overnight. We sampled individual flower buds of young main inflorescences but avoided the first and second flowers of the inflorescence because these flowers tend to show developmentally immature morphologies. We collected flower buds with mature pollen but before the anthers were opened (flower stage 13), and added 30 µl of 5% Tween 20 (Sigma-Aldrich, St. Louis, MO, USA) to each tube. The tubes were sonicated using a Bioruptor (Diagenode, Seraing, Belgium) in high power mode with 10 cycles of sonication-ON for 30 s and sonication-OFF for 30 s so that the pollen grains were released from the anther sacs. After a short centrifugation and vortexing, 10 µl of the solution was mounted on a Neubauer slide. We took three images per sample using a light microscope. The number of pollen grains per image was counted using the particle counting implemented in ImageJ (http://imagej.nih.gov/ij/) and in Fiji (http://fiji.sc/Fiji). We then estimated the total pollen number per flower based on the image size and the total volume. Ovule numbers were counted by dissecting young siliques under a dissecting microscope.

Because of limited chamber space, we split the plants into two batches. The two batches were treated under the same conditions in the same chambers, but at different times (see Extended Data Table 2 for details). We controlled this potential batch effect by setting equal medians and standard deviations for the two batches.

Sometimes there were no or very few pollen grains per image. This was mainly in situations where anthers did not open. To eliminate these artefacts, we discarded the flowers with pollen counts of less than 10 per image. We confirmed that such extremely low pollen numbers did not appear in specific accessions, indicating that this is not heritable.

### Plant materials and growth conditions for functional analyses

For the functional analyses, we mainly used *Arabidopsis thaliana* wild types and mutants of the Col-0 and Bor-4 accessions. The T-DNA lines SALK_064854/N666274 (*rdp1-1*) from the Salk collection^33^, GK-879G09/N484369 (*rdp1-2*) from GABI-Kat lines^34^, and GT_5_108298/N180401 from JIC Gene Traps^35^ were obtained from the European *Arabidopsis* Stock Centre^36^.

The T-DNA insertion in each line was confirmed using PCR with the primers listed in Table S7 as described at http://signal.salk.edu/tdnaprimers.2.html. DNA was extracted from young leaves using the CTAB method.

We generated single nucleotide insertion/deletion lines in the Col-0 and Bor-4 accessions using the FAST-CRISPR-Cas9 construct (see the section “CRISPR mutant”) and designated them *rdp1-3* and *rdp1-4* (in Col-0), and *rdp1-5* (in Bor-4).

*Arabidopsis* seeds were sown on soil mixed with the insecticide ActaraG (Syngenta Agro, Switzerland) and stratified for 3–4 days at 4 °C in the dark. The plants were grown under 16 h of light at 22 °C, and 8 h of dark at 20 °C, with weekly treatments of insecticide (Kendo Gold, Syngenta Agro) unless noted otherwise (for GWAS, see above).

### Statistical analysis

Unless otherwise stated, statistical and population genetic analyses were performed using the statistical software R^37^. In bar plots, bars indicate the median, boxes indicate the interquartile range, and whiskers extend to the most extreme data point that is no more than 1.5 times the interquartile range from the box, with outliers shown by dots.

### Histological analysis of anthers

Inflorescences were fixed with formaldehyde:acetic acid:70% ethanol = 1:1:18 (FAA), dehydrated and embedded in Technovit 7100 according to the manufacturer’s instructions (Heraeus Kulzer GmbH, Wehrheim, Germany) for the histological analysis. Five-micrometer sections were cut with a microtome (RM2145, Leica, Germany) and stained with toluidine blue before observation under a Leica microscope (DM5000, Leica) equipped with a black-and-white camera (DFC345, Leica).

### Correlations and overlap with published GWAS results and climatic data

To examine whether pollen and ovule numbers were correlated with any of the other 107 published phenotypes^17^ or with climate and geographic variables^18^, Pearson’s correlation coefficients were calculated (Extended Data Tables 4 and 5). We also surveyed whether there were any SNPs significantly associated with climate variables in the 10-kb window including the SNP of the highest GWAS score of pollen number (Chr1:8,850,000– 8,860,000). The significance was based on the genome-wide empirical *P*-values^18^. Focusing on the same 10-kb window, we also surveyed whether there were significant SNPs (*P* < 10^−5^; minor allele frequency > 0.1) for the 107 published phenotypes in the GWAS. The correlation of pollen number and the S-haplogroups^19^ was also tested.

### SNP imputation

To perform GWAS, we generated a set of dense SNP markers that overlapped with the phenotyped accessions using imputation^38^. First, we constructed a reference set of 186 haplotypes from resequencing data^39,40^. MaCH version 1.0.16.c^41^ was then used to impute nongenotyped SNPs for 1311 accessions in 250 k SNP array data^22^. The command line used for each overlapping bin was:

mach1--dosage --greedy -r -d [sample_bin].dat -p [sample_bin].ped -s [ref_bin].snps -h [ref_bin].haplos --prefix [output_prefix]

The output was then merged and converted into the homozygous SNP dataset.

### Genome-wide association study (GWAS)

GWAS was performed to identify loci associated with pollen number variation in the 144 natural accessions. We also performed GWAS for ovule number and for the 107 published phenotypes^17^ using the same SNP set as for pollen number. The median values of pollen number were used to represent each accession. We used the log-transformed values of pollen number for GWAS, as they did not deviate significantly from a normal distribution (Shapiro–Wilk normality test: *P* = 0.438). To deal with confounding as a result of population structure, we employed the mixed model implemented in the software Mixmogam, in which a genome-wide kinship matrix was incorporated as a random effect^31^. By using Mixmogam, we also generated a quantile–quantile (Q–Q) plot, which shows the relationship between the observed and expected negative logarithm of *P*-values (Fig. 1h). For the Manhattan plot (Fig. 1f, g), we removed SNPs with minor allele frequencies < 0.15, leaving 1,115,178 SNPs overall.

### RNA extraction, cDNA synthesis, and expression analysis

For gene expression analysis, total RNA was isolated using the ZR Plant RNA MiniPrep (Zymo Research, Irvine, CA) according to the manufacturer’s instructions. Total RNA was treated with a DNA-free DNA removal kit (Thermo Fisher Scientific, Waltham, MA) and cDNA was synthesized using a High-Capacity RNA-to-cDNA kit (Thermo Fisher Scientific). For quantitative expression analysis, cDNA was used with SYBR Select Master Mix (Thermo Fisher Scientific) and gene-specific primers (Extended Data Table 7). Data were collected using the StepOnePlus Real-Time PCR System (Thermo Fisher Scientific) in accordance with the instruction manual. Expression levels were normalized using *EF1-α* (*AT5G60390*), which was used as an internal control.

### Measuring pollen number and size with a cell counter

To expedite pollen number counting, we established a rapid method using a cell counter (CASY TT, OMNI Life Science GmbH, Germany). We found that the pollen numbers of flowers on side inflorescences and side branches of the main inflorescence were similar, but those of flowers on the main inflorescence were higher (Extended Data Fig. 12). To obtain a large number of replicates, we sampled flowers from the former two positions. We sampled during the first 3 weeks of flowering and excluded the first and second flowers on each branch; this yielded up to 40 flowers per individual. Collecting, suspending, and sonicating of flowers for GWAS were conducted as described above. All pollen solutions were suspended in 10 ml of CASYton (OMNI Life Science GmbH), and pollen numbers were counted with a CASY TT cell counter as described previously^42^. Three 400 µl aliquots of each pollen solution were counted. We counted particles within a size range of 12.5–25 µm (estimated diameter) as pollen. Samples with a clear peak at 7.5–12.5 µm were discarded as broken or unhealthy samples.

### Transgenic experiment

The *RDP1* genomic sequence was amplified by PCR using Phusion High Fidelity PCR polymerase (New England Biolabs, Beverly, MA). We used the primers 3550_At1g25260F1 (5´-TTTCTCCCCACATTTCTC-3´) and 3551_At1g25260R1 (5´-GTTTAAAATGAGAGAACCCG-3´) to amplify the full length of *RDP1* and the surrounding genomic sequence that may contain promoter and terminator regions (ca. 4.3 kbp, positions 8,857,725 to 8,853,420 on chromosome 1). The *RDP1* promoter sequence was amplified by PCR using the following primers: 3550_At1g25260F1 (5´-TTTCTCCCCACATTTCTC-3´) and 4155_At1g25260R (5´-AGCTGAGGTTTCAAATGTTGTATG-3´). These sequences were cloned into a pCR8 vector using the pCR8/GW/TOPO TA Cloning kit (Thermo Fisher Scientific). pCR8-RDP1 complete sequences were cloned into a pFAST-G01 vector^43^, and the pCR8-RDP1 promoter vector was cloned into a pGWB3 vector^44^ using Gateway LR clonase II (Thermo Fisher Scientific). We designated these constructs pFAST-RDP1 and pGWB3-RDP1pro, respectively. Plants were grown at 22 °C under a 16 h light/8 h dark cycle until transformation via *Agrobacterium tumefaciens* strain GV3101 using the floral-dip method^45^. pFAST-RDP1 was transformed into *rdp1-1* homozygous mutants. Transformed seeds were selected using a fluorescence stereomicroscope with a GFP filter (Olympus SZX12, Japan). pGWB3-RDP1pro was transformed to Col-0. Seedlings carrying pGWB3-RDP1pro were selected on 1/2 Murashige and Skoog medium supplemented with 50 µg/ml kanamycin and 50 µg/ml hygromycin.

### CRISPR mutants

To generate *RDP1* frameshift mutants, the CRISPR-Cas9 system was used as previously described^24^. A 20-nucleotide target sequence (5´-GCAGAAGATGAACTCCGTTC-3´) was selected using the CRISPR-P tool^46^. The target sequence was subcloned into a pTTK194 vector (pUC19 U6.26pro::sgRNA). Then, the *Hin*dIII sgRNA fragment was subcloned into a pTTK182 (pFAST-R02 35Spro::Cas9) vector or pTTK227 (pFAST-R01 RPS5Apro::Cas9). For plant transformation, the binary vector was first introduced into *Agrobacterium tumefaciens* (GV3101) and then into *A. thaliana* by the floral-dip method.

### Counting the microspore mother cells

Flower buds were collected and fixed in 3:1 (vol:vol) ethanol:acetic acid for 16 h. The fixed flower buds were rehydrated with 99.5%, 70%, and 50% ethanol for 30 min each.Flower buds were transferred to 1 N NaOH for 3 h and then stained with aniline blue solution (0.1% aniline blue, 100 mM K_3_PO_4_) for 3 h. Anthers at the microspore mother cell stage were dissected under a microscope with UV illumination (DM5000, Leica, Germany). Z-stack images were obtained with a confocal microscope (SP5, Leica).

### *In situ* hybridization

Flower buds were fixed with 3.7% formaldehyde, 5% acetic acid, 50% ethanol (FAA) and dehydrated through an ethanol series. Fixed samples were embedded in paraplast using an embedding machine (ASP200, Leica). *RDP1* cDNA was PCR-amplified using the primer pair (At1g25260g2F; 5´-tgcctaatcaaagcgagtagacc-3´ and At1g25260gR; 5´-cagagcaagttcagcttgaaagtagc-3´) and cloned into pCR4-TOPO (Thermo Fisher Scientific) vector. Cloned cDNA was used as a template for *in vitro* transcription using a MAXIscript T7 labeling kit (Thermo Fisher Scientific). *In situ* mRNA hybridization was performed as described previously^32^.

### GUS assay

Plant samples were incubated in 90% acetone for 20 min at room temperature, washed with 50 mM phosphate buffer containing 0.1% Triton X-100, 2 mM potassium ferrocyanide, and 2 mM potassium ferricyanide, and incubated in the same buffer supplemented with 1 mg/ml X-Gluc for 5 h at 37 °C^47^.

### Phylogenetic analysis

Multiple sequence alignment was performed using ClustalW implemented in the CLC Workbench (version 7.7). A phylogenetic tree was generated by the neighbour-joining distance algorithm using the aligned region (amino acid positions 44–153) and a bootstrap value of 1,000. Yvh1 of *Saccharomyces cerevisiae* was used as an outgroup. Accession numbers are listed below in the Accession Numbers section.

### Selection scan

For the selection scan, we used the imputed SNP dataset that was also used for GWAS. We used 298 accessions (Extended Data Table 1), covering all the accessions used for our GWAS of pollen and ovule numbers and the GWAS of 107 phenotypes reported by Atwell *et al*.^17,22,48^. We used the iHS statistic for the selection scan; this statistic compares the extended haplotype homozygosity (EHH) between two alleles by controlling for the allele frequency of each SNP^20^. The R library rehh was used to calculate the iHS statistics^49^. The *Arabidopsis lyrata* reference genome was used to infer the ancestral state for each SNP^50^.We first calculated the iHS for each SNP. Then, as in Horton *et al*.^22^, we split the genome into 10-kb windows and used the maximum score from the iHS scan for each window as the test statistic. Empirical *P*-values were calculated for all windows and for all SNPs.

We then asked whether the GWAS-associated windows were enriched in the extreme tails of the iHS statistic. Enrichment analysis was performed across the 10-kb windows. To examine whether enrichment of the iHS was commonly observed in GWAS peaks, we also performed this analysis for the GWAS results for the other 107 publicly available phenotypes^17^ and compared them with the GWAS of pollen and ovule numbers. Four thresholds of empirical *P*-value tails were considered: 10%, 5%, 2.5%, and 1%. GWAS SNPs were considered if *P*-values were smaller than 10^−4^ and the minor allele frequency was > 0.1.

The statistical significance of the fold enrichment was addressed based on permutation tests that preserve the linkage disequilibrium structure in the data. A set of windows was resampled for each permutation, preserving the relative positions of the windows, but shifting them by a randomly chosen uniformly drawn number of windows for each permutation. A similar method of permutation has been used in several population genomic studies^17,22,48^. Permutation was performed 1,000 times.

### Seed and fruit number determinations

All the fruits on the main inflorescence were counted more than 65 days after germination. Unopened flower buds at the tip were not counted as fruits. Fruits at the intermediate positions were opened and their number of seeds was scored. Immature seeds were also counted as seeds.

### Rosette size determination using Fiji (ImageJ)

We took images of plants that included a ruler at 3 weeks after germination. A minimum circumscribed circle was drawn manually on the picture using Fiji; then, the area was measured and transformed depending on the scale.

### Accession numbers

Sequence data from this article can be found in The Arabidopsis Information Resource and GenBank (National Center for Biotechnology Information) databases under the following accession numbers: RDP1/Mrt4: *Arabidopsis thaliana*; AT1G25260 (Col-0), LC164158 (Mz-0), LC164159 (Bor-4), *Arabidopsis lyrata*; LC164160, *Arabidopsis halleri*; LC164161, *Capsella rubella*; XP006304175, *Theobroma cacao*; XP0074155, *Populus trichocarpa*; XP002299012, *Vitis vinifera*; XP002276595, *Solanum lycopersicum*; XP004230519, *Prunus persica*; XP007223904, *Lotus japonicus*; AFK33945, *Glycine max*; XP006584472, *Medicago truncatula*; XP003593096, *Oryza sativa*; NP001065532, *Zea mays*; DAA39345, *Physcomitrella patens*; XP001771523, *Acanthamoeba castellanii*; XP004351551, *Caenorhabditis elegans*; NP495470, *Latimeria chalumnae*; XP005986121, *Homo sapiens*; AF173378_1, *Astyanax mexicanus*; XP007251523, *Drosophila virilis*; XP002050865, *Drosophila melanogaster*; NP610554, *Saccharomyces cerevisiae*; gi330443667, P0 protein: *Arabidopsis thaliana*; AT3G09200, AT3G11250, AT2G40010, *Arabidopsis lyrata*; EFH58973, *Arabidopsis halleri*; LC164162, LC164163, *Drosophila melanogaster*; NP524211, *Homo sapiens*; NP000993, *Caenorhabditis elegans*; NP492766, *Saccharomyces cerevisiae*; NP013444, Yvh1: *Saccharomyces cerevisiae*; NP01229.

## Reference

1. Harvey, P. H. & May, R. M. Out for the sperm count. Nature 337, 508–509 (1989).

2. Shimizu, K. K. & Tsuchimatsu, T. Evolution of selfing: recurrent patterns in molecular adaptation. Annu. Rev. Ecol. Evol. Syst. 46, 593–622 (2015).

3. Birkhead, T. R. & Møller, A. P. Sperm Competition and Sexual Selection (Academic Press, 1998).

4. Merzenich, H., Zeeb, H. & Blettner, M. Decreasing sperm quality: a global problem? BMC Public Health 10, 24 (2010).

5. Aitken, R. J. Falling sperm counts twenty years on: where are we now? Asian J. Androl. 15, 204–207 (2013).

6. Czeizel, A. E. & Rothman, K. J. Does relaxed reproductive selection explain the decline in male reproductive health? A new hypothesis. Epidemiology 13, 113–114 (2002).

7. Willis, J. H. The contribution of male-sterility mutations to inbreeding depression in Mimulus guttatus. Heredity 83, 337–346 (1999).

8. Fu, W. & Akey, J. M. Selection and adaptation in the human genome. Annu. Rev. Genomics Hum. Genet. 14, 467–489 (2013).

9. Oka, H.-I. & Morishima, H. Variations in the breeding systems of a wild rice, Oryza perennis. Evolution 21, 249–258 (1967).

10. Barrett, S. C. H. The evolution of plant sexual diversity. Nat. Rev. Genet. 3, 274–284 (2002).

11. Darwin, C. The Effects of Cross and Self Fertilisation in the Vegetable Kingdom (John Murray, 1876).

12. Sicard, A. & Lenhard, M. The selfing syndrome: a model for studying the genetic and evolutionary basis of morphological adaptation in plants. Ann. Bot. 107, 1433–1443 (2011).

13. Kosova, G., Scott, N. M., Niederberger, C., Prins, G. S. & Ober, C. Genome-wide association study identifies candidate genes for male fertility traits in humans. Am. J. Hum. Genet. 90, 950–961 (2012).

14. Tsuchimatsu, T. et al. Evolution of self-compatibility in Arabidopsis by a mutation in the male specificity gene. Nature 464, 1342–1346 (2010).

15. Cruden, R. W. Pollen grains: why so many? Plant Syst. Evol. 222, 143–165 (2000).

16. Charnov, E. L. The Theory of Sex Allocation (Princeton University Press, 1982).

17. Atwell, S. et al. Genome-wide association study of 107 phenotypes in Arabidopsis thaliana inbred lines. Nature 465, 627–631 (2010).

18. Hancock, A. M. et al. Adaptation to climate across the Arabidopsis thaliana genome. Science 334, 83–86 (2011).

19. Shimizu, K. K., Shimizu-Inatsugi, R., Tsuchimatsu, T. & Purugganan, M. D. Independent origins of self-compatibility in Arabidopsis thaliana. Mol. Ecol. 17, 704–714 (2008).

20. Voight, B. F., Kudaravalli, S., Wen, X. & Pritchard, J. K. A map of recent positive selection in the human genome. PLoS Biol. 4, e72 (2006).

21. Tishkoff, S. A. et al. Haplotype diversity and linkage disequilibrium at human G6PD: recent origin of alleles that confer malarial resistance. Science 293, 455–462 (2001).

22. Horton, M. W. et al. Genome-wide patterns of genetic variation in worldwide Arabidopsis thaliana accessions from the RegMap panel. Nat. Genet. 44, 212–216 (2012).

23. Jinek, M. et al. A programmable dual-RNA-guided DNA endonuclease in adaptive bacterial immunity. Science 337, 816–821 (2012).

24. Tsutsui, H. & Higashiyama, T. pKAMA-ITACHI vectors for highly efficient CRISPR/Cas9-mediated gene knockout in Arabidopsis thaliana. Plant Cell Physiol. 58, 46–56 (2017).

25. Rodríguez-Mateos, M. et al. Role and dynamics of the ribosomal protein P0 and its related trans-acting factor Mrt4 during ribosome assembly in Saccharomyces cerevisiae. Nucl. Acids Res. 37, 7519–7532 (2009).

26. Bhavsar, R. B., Makley, L. N. & Tsonis, P. A. The other lives of ribosomal proteins. Hum. Genomics 4, 327–344 (2010).

27. Sanders, P. M. et al. Anther developmental defects in Arabidopsis thaliana male-sterile mutants. Sex. Plant Reprod. 11, 297–322 (1999).

28. Breuninger, H. & Lenhard, M. Control of tissue and organ growth in plants. Curr. Top. Dev. Biol. 91, 185–220 (2010).

29. Turner, T. L. Fine-mapping natural alleles: quantitative complementation to the rescue. Mol. Ecol. 23, 2377–2382 (2014).

30. Stern, D. L. A role of Ultrabithorax in morphological differences between Drosophila species. Nature 396, 463–466 (1998).

## Reference

31. Segura, V. et al. An efficient multi-locus mixed-model approach for genome-wide association studies in structured populations. Nat. Genet. 44, 825–830 (2012).

32. Peterson, K. M. et al. Arabidopsis homeodomain-leucine zipper IV proteins promote stomatal development and ectopically induce stomata beyond the epidermis. Development 140, 1924–1935 (2013).

33. Alonso, J. M. et al. Genome-wide insertional mutagenesis of Arabidopsis thaliana. Science 301, 653–657 (2003).

34. Kleinboelting, N., Huep, G., Kloetgen, A., Viehoever, P. & Weisshaar, B. GABI-Kat SimpleSearch: new features of the Arabidopsis thaliana T-DNA mutant database. Nucleic Acids Res. 40, D1211–D1215 (2012).

35. Sundaresan, V. et al. Patterns of gene action in plant development revealed by enhancer trap and gene trap transposable elements. Genes Dev. 9, 1797–1810 (1995).

36. Scholl, R. L., May, S. T. & Ware, D. H. Seed and molecular resources for Arabidopsis. Plant Physiol. 124, 1477–1480 (2000).

37. R Core Team. R foundation for statistical computing. Vienna Austria (2013).

38. The 1001 Genomes Consortium. 1,135 genomes reveal the global pattern of polymorphism in Arabidopsis thaliana. Cell 166, 481–491 (2016).

39. Cao, J. et al. Whole-genome sequencing of multiple Arabidopsis thaliana populations. Nat. Genet. 43, 956–965 (2011).

40. Long, Q. et al. Massive genomic variation and strong selection in Arabidopsis thaliana lines from Sweden. Nat. Genet. 45, 884–890 (2013).

41. Li, Y., Willer, C. J., Ding, J., Scheet, P. & Abecasis, G. R. MaCH: using sequence and genotype data to estimate haplotypes and unobserved genotypes. Genet. Epidemiol. 34, 816–834 (2010).

42. Tedder, A. et al. Evolution of the selfing syndrome in Arabis alpina (Brassicaceae). PLoS One 10, e0126618 (2015).

43. Shimada, T. L., Shimada, T. & Hara-Nishimura, I. A rapid and non-destructive screenable marker, FAST, for identifying transformed seeds of Arabidopsis thaliana. Plant J. 61, 519–528 (2010).

44. Nakagawa, T. et al. Improved gateway binary vectors: high-performance vectors for creation of fusion constructs in transgenic analysis of plants. Biosci. Biotechnol. Biochem. 71, 2095–2100 (2007).

45. Clough, S. J. & Bent, A. F. Floral dip: a simplified method forAgrobacterium-mediated transformation ofArabidopsis thaliana. Plant J. 16, 735–743 (1998).

46. Lei, Y. et al. CRISPR-P: A web tool for synthetic single-guide RNA design of CRISPR-system in plants. Mol. Plant 7, 1494–1496 (2014).

47. Vielle-Calzada, J. P., Baskar, R. & Grossniklaus, U. Delayed activation of the paternal genome during seed development. Nature 404, 91–94 (2000).

48. Nordborg, M. et al. The pattern of polymorphism in Arabidopsis thaliana. PLoS Biol. 3, 1289–1299 (2005).

49. Gautier, M. & Vitalis, R. rehh: an R package to detect footprints of selection in genome-wide SNP data from haplotype structure. Bioinformatics 28, 1176–1177 (2012).

50. Hu, T. T. et al. The Arabidopsis lyrata genome sequence and the basis of rapid genome size change. Nat. Genet. 43, 476–481 (2011).

